# CluSeek: Bioinformatics Tool to Identify and Analyze Gene Clusters

**DOI:** 10.1101/2025.09.16.676505

**Authors:** Ondrej Hrebicek, Stanislav Kadlcik, Lucie Najmanova, Jiri Janata, Jana Kamanova, Lada Hanzlikova, Marketa Koberska, Vojtech Kovarovic, Zdenek Kamenik

## Abstract

Gene clusters are key structural and functional units that encode diverse phenotypes in genomes, from metabolism to pathogenesis. As genome sequencing expands, tools for systematic exploration of this growing data are increasingly needed. We present CluSeek, an open-source, Python-based platform for discovering, visualizing, and analyzing gene clusters across all GenBank data. Unlike existing tools, CluSeek does not rely on predefined cluster types or reference libraries but enables mining of any gene neighborhoods containing colocalized homologs of user-specified genes. It features an intuitive graphical interface suitable for non-bioinformaticians and is freely available at https://cluseek.com. We demonstrate the versatility of Cluseek in two distinct case studies: (i) mining of specialized metabolites, where CluSeek uncovered over 16 new classes containing the bioactivity-enhancing 4-alkyl-L-proline moiety, previously known in only three Golden Era antibiotic classes; and (ii) analysis of type III secretion systems present in *Bordetella* species, revealing previously unrecognized taxonomic distribution, and genetic variants, including gene multiplications and novel components with potential functional significance.

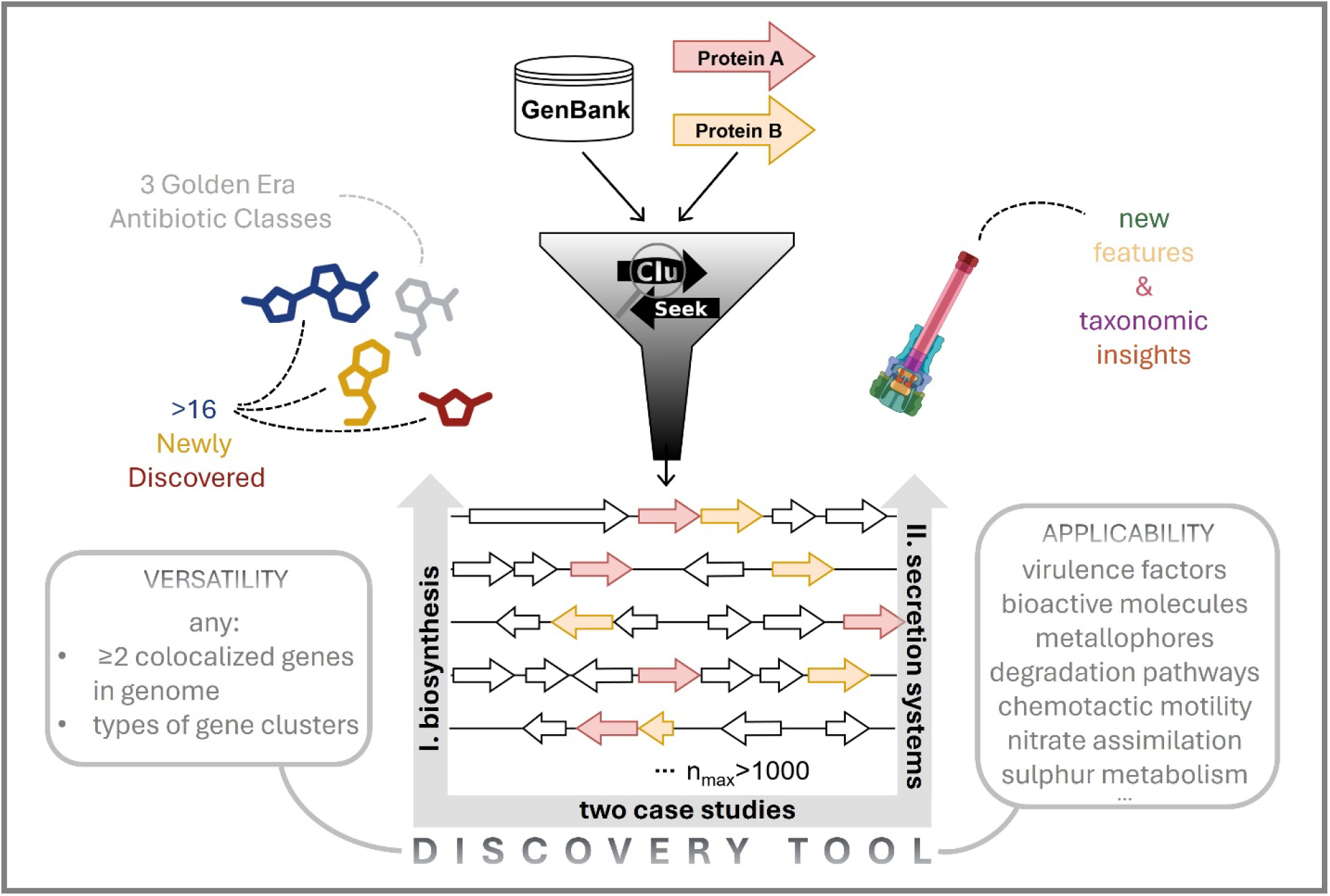

## INTRODUCTION

In bacteria, archaea, and fungi, genes encoding complex phenotypes or processes are typically arranged into gene clusters, including those for antibiotic, signaling and iron-acquisition molecule biosynthesis; two-component regulatory systems; chemotactic motility; nitrate assimilation and sulfur metabolism machineries; and secretion systems^1–5^. Such gene organization has two major consequences. First, it facilitates regulation of the respective process via gene co-expression. Second, tightly preserved clusters allow for efficient horizontal gene transfer between organisms regardless of their phylogenetic relatedness. This phenomenon represents an important evolutionary driving force which not only spreads functional gene units into new hosts but also enables modular grouping of simple gene clusters (subclusters) into more complex ones^1^. It results in vast diversity of various aspects of the microbial world, be it the pool of bioactive metabolites produced by microorganisms or the variety of molecular mechanisms of pathogens to hijack host cells.

The past two decades have seen the increasing accumulation of a vast amount of genomic data in public databases, which encompasses both individual laboratory research topics, as well as high throughput sequencing of public microbial strain collections. Combined, this data represents a largely unexplored treasure trove of yet-undescribed gene (sub)clusters and it offers a great opportunity for targeted genome mining to discover various novel or unusual phenomena^6,7^. However, the ever-increasing volume of available data necessitates the development of more sophisticated genome exploring and mining tools which allow the user to access and analyze this data in a time-efficient manner. Those widely used tools, e.g. antiSMASH for metabolite biosynthesis^8^ or SecReT6 for secretion systems^9^, identify gene clusters based on curated libraries comprising a limited set of previously characterized examples and/or general rules derived from well-studied, canonical cases. This dependence on the available knowledge also means that these tools are specialized in a certain area of research. As such they readily provide an overview of gene clusters (limited to the specialization of the tool) in a restricted amount of genomic data uploaded by the user. However, these library- and rule-based tools predominantly reveal already known gene clusters and their variants, while unique, unusual or yet-undiscovered types may be overlooked. To address this, we developed a Python-based genome mining tool CluSeek, ‘Cluster Seeker’, which searches for colocalized homologs of any marker genes/proteins (markers) within the GenBank database. The tool retrieves and analyzes gene clusters which contain the user-defined marker genes with no regard to the size, overall gene composition or function of the cluster. CluSeek is publicly accessible, it can be easily launched and owing to integrated graphical user interface (GUI), it can be operated by users with no IT or bioinformatics background. We demonstrate its applicability and versatility using two distinct case studies: (i) biosynthesis of specialized metabolites by Gram-positive Actinomycetes (ii) secretion systems as virulence factors of Gram-negative pathogens.

## RESULTS AND DISCUSSION

### CluSeek Workflow

CluSeek is a versatile genome mining tool, which (i) searches for gene clusters which share homologs of two or more genes of user interest, (ii) scans through large genomic data deposited in GenBank, (iii) is independent of libraries of known gene clusters, (iv) does not rely on pre-defined gene cluster-detecting rules, (v) is not restricted to a specific research area.

Fig. 1A illustrates the CluSeek processing pipeline. It begins with a BLASTp^10^ search of user-selected markers against the non-redundant (nr) protein database, which can be performed in two ways: (i) by providing amino acid sequences in FASTA format, allowing CluSeek to run the BLASTp search against the nr database directly through its GUI; or (ii) by running BLASTp externally (e.g., via the NCBI website - ensuring that the nr database is selected) and uploading the resulting XML file. BLASTp hits can be optionally filtered using interactive histograms to restrict the results to homologs with a defined level of sequence similarity (based on e-value or sequence identity). To determine the positions of the identified marker homologs, CluSeek uses the NCBI Identical Protein Group (IPG) resource, which provides information on the GenBank sequences, in which the homologs are found, and their exact position. Based on this information, CluSeek identifies genome sequences where the markers are colocalized within a user-defined (or default) genomic distance. Next, CluSeek retrieves the GenBank records of genes neighboring the markers from the Nuccore database^11^, hereinafter referred to as gene clusters. All proteins in the region are then assigned into protein subgroups based on sequence similarity using the Diamond aligner^12^. These subgroups are further refined into larger, less homogeneous groups of homologous proteins via community detection on a local alignment-based network using the NetworkX library^13^ (see *Methods*, section *Gene Cluster Visualization and Protein Grouping* for details). This enables the identification of homologous proteins across all retrieved gene clusters. As the original subgroups are retained and accessible to the user, CluSeek supports downstream analyses at both the protein group and subgroup levels, enhancing the tool’s versatility. Finally, CluSeek visualizes the gene clusters and orders them by pairwise similarity, calculated using the Jaccard index of shared protein groups, ensuring that similar gene clusters are displayed in proximity.

**Fig. 1.**
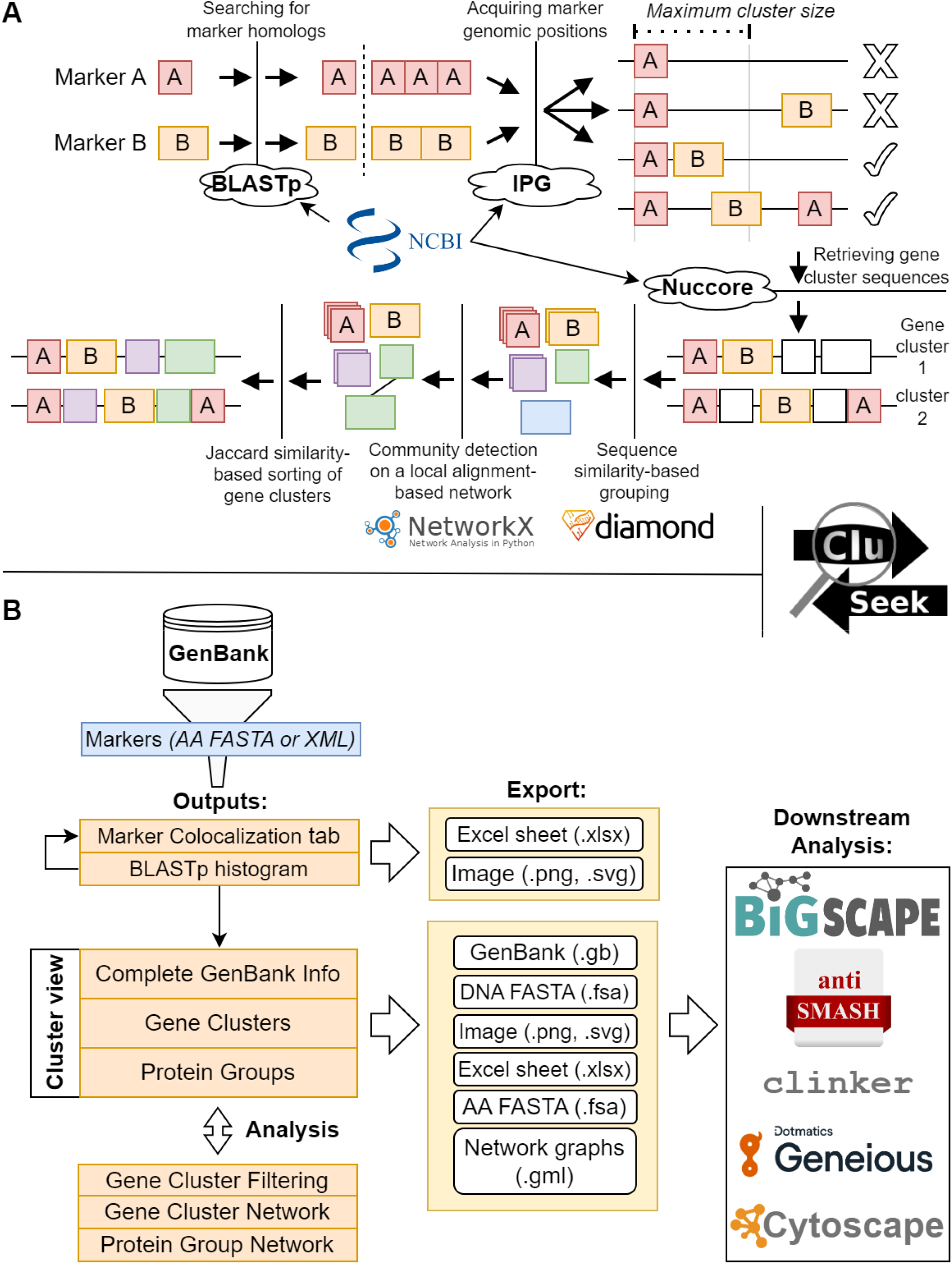
Overview of CluSeek. **A. Workflow for two marker-based gene cluster detection**. Marker homologs are identified in GenBank via BLASTp against the nr database and can be optionally filtered based on sequence similarity; sequences containing the markers within a user-defined distance are selected using IPG resource; gene neighborhoods are retrieved from GenBank via the Nuccore database; all encoded proteins are grouped in two steps: by sequence similarity using Diamond, and by community detection on a local alignment-based network graph using NetworkX; visualized gene clusters are ordered by the number of shared protein groups. **B. Outputs and GUI features** CluSeek retrieves and visualizes gene clusters from GenBank based on user-selected markers. It provides: (i) a table of species with colocalized marker homologs; (ii) a histogram with marker homologs identified by BLASTp; and (iii) a gene cluster view as the main output showing visualized gene clusters with full GenBank information and corresponding protein groups. The GUI supports analyses by taxon, gene content, or similarity networks. The outputs can be exported for downstream use in various third-party tools.

CluSeek provides an intuitive GUI that guides users through the workflow, visualizes its outputs, and enables their export in multiple formats (Fig. 1B). The GUI consists of two sections: (1) the *Markers and Colocalization* tab, and (2) the *Cluster View* tab. To help users initiate their analysis and navigate the interface, we provide a set of video tutorials available on our YouTube channel https://www.youtube.com/@CluSeek-p6u.

The *Markers and Colocalization* tab allows users to select the markers and perform colocalization analysis. Its results are displayed in two sub-tabs: *Marker Homologs* and *Colocalization*. The *Marker Homologs* sub-tab summarizes the BLASTp results, including a histogram with the distribution of homologs for each marker based on e-value or sequence identity. While optional, this feature helps users assess the specificity of their selected markers - an important step, as a high number of homologs (e.g., exceeding NCBI’s *max target sequences* return limit) may hinder or bias downstream analysis. Based on this output, users may refine their marker selection, restrict the BLASTp searches to specific taxa, or select only a subset of homologs for re-colocalization. The *Colocalization* sub-tab lists organisms in which marker homologs occur within a user-defined genomic distance, identifying those that contain the targeted gene clusters.

The *Cluster View* tab consists of four sub-tabs, including CluSeek’s primary output represented by an interactive, customizable visualization of identified gene clusters (*Gene Clusters* sub-tab). Here, users can toggle between fixed and proportional gene sizes, customize gene colors and labels, align clusters to a gene of interest, and access detailed information from GenBank, including accession numbers, sequences, and functional annotations, with interactive links to the corresponding GenBank entries (*Info* sub-tab). CluSeek also supports filtering of gene clusters based on contained genes (e.g., presence of a specific gene and its homologs; *Filtering* sub-tab) or by taxa (*Gene Clusters* sub-tab). Users can inspect individual protein groups (*Protein Groups* sub-tab), and via protein group networks (*Info* sub-tab), they can merge selected protein groups, split them into sequence similarity-based subgroups, or adjust groupings based on external information. Since CluSeek can process large numbers of gene clusters – potentially overwhelming to explore individually – it also offers the option to analyze gene cluster diversity via a gene cluster network. This network groups clusters based on shared genes, providing an intuitive overview of their relationships. The resulting network graph (*Export* sub-tab) can be exported for visualization and interpretation in Cytoscape^14^.

Finally, CluSeek outputs are compatible with third-party tools such as antiSMASH (for further characterization and comparison to known biosynthetic gene clusters) or Geneious (e.g., for primer design in heterologous expression workflows).

### Case study 1 - Biosynthesis

Natural products are highly valued for their various medically important bioactivity and remarkable structural diversity, which is achieved by combining gene subclusters representing individual metabolite building blocks and gene gain, loss, or modification that determine metabolite tailoring or means of building blocks assembly^1^. In this case study, we employed CluSeek for genome mining of metabolites containing an unusual structural building block, 4-alkyl-L-proline. This building block is biosynthesized from L-tyrosine by five or six enzymes named Apd1-6^15^, which are encoded within the *apd* subcluster, and it is then incorporated into three known classes of bioactive metabolites produced by Actinomycetota: lincomycin^16,17^, hormaomycin^18^, and pyrrolobenzodiazepines^19^ (Fig. 2A). While 4-alkyl-L-proline is the only common feature among these metabolites, they differ in both overall structure and bioactivity. Lincomycin is a clinically used antimicrobial composed of a 4-alkyl-L-proline and an amino thiosaccharide building block that targets Gram-positive pathogens by binding to their 50S ribosomal subunit^20,21^. Hormaomycin is a complex cyclic peptidic lactone composed of eight amino acid residues, seven of which are non-proteinogenic. It is considered a signaling molecule with a narrow-spectrum antimicrobial activity^22^. Pyrrolobenzodiazepines are a large class of tricyclic metabolites that act as antitumor agents by binding to a minor groove of DNA^19^. The 4-alkyl-L-proline structural motif within these metabolites is crucial, as it enhances the bioactivity of functionally distinct lincomycin and pyrrolobenzodiazepines compared to their analogs with simple L-proline instead^23,24^.

**Fig. 2.**
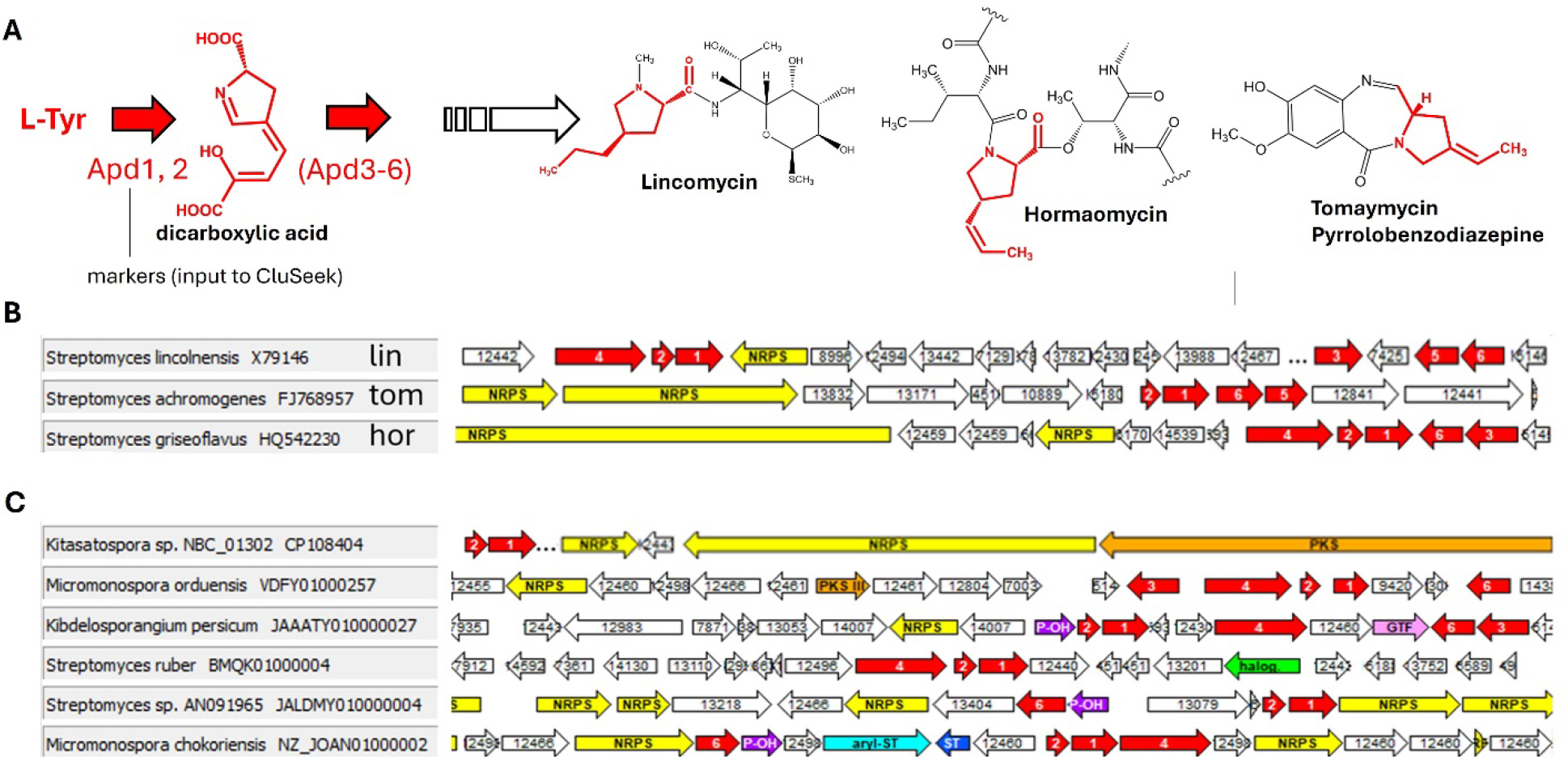
4-alkyl-L-proline-containing microbial metabolites. Biosynthesis, structures (A), and BGCs (B) representing all known classes of these metabolites: lincomycin (lin), hormaomycin (hor) and pyrrolobenzodiazepines represented by tomaymycin (tom); (C) Examples of unique putative biosynthetic gene clusters newly identified by CluSeek (the complete CluSeek output is depicted in Supplementary Figure S1). The clusters contain the *apd* subcluster genes (in red), but are distinct from those clusters of known metabolites; genes encoding Apd1-6 proteins (labeled 1-6) - in red; sulfotransferase (ST) - in blue; halogenase - in green, polyketide synthase (PKS), type III PKS (PKS III) - in orange; aryl-sulfotransferase (aryl-ST) - in turquoise; L-proline 3-hydroxylase (P-OH) - in violet; glycosyltransferase (GTF) - in pink; non ribosomal peptide synthase (NRPS) - in yellow; five-digit numbers are randomly assigned by CluSeek and can be replaced with gene names; identical numbers mark homologous genes. Three dots indicate omitted genes from long gene clusters. Gene cluster borders are not shown; only selected gene regions are displayed.

To expand our limited knowledge of only three classes of metabolites with a 4-alkyl-L-proline building unit, we used Apd1 and Apd2 proteins, which convert L-tyrosine into a 4-alkyl-L-proline biosynthesis-specific dicarboxylic acid^15,25^, as markers (see Fig. 2A). Using CluSeek on the GenBank database, we mined 508 gene clusters that contain genes of the *apd* subcluster (Supplementary Fig. S1). Out of these clusters, only seven had been previously published as biosynthetic gene clusters (BGCs) associated with a metabolite of known structure, i.e., lincomycin^26^, hormaomycin^18^ and five pyrrolobenzodiazepines: anthramycin^27^, tomaymycin^28^, sibiromycin^29^, porothramycin^30^ and limazepine^31^ (representative examples are presented in Fig. 2B). Out of the newly mined gene clusters, 191 (38%) are related to known gene clusters. Specifically, 5 are likely related to lincomycin, 9 to hormaomycin, and 177 to pyrrolobenzodiazepines or their structural analogs (see Fig. 3A). However, we were particularly intrigued by the discovery of the 317 (62%) additional gene clusters that were markedly distinct from known BGCs. These represent candidate BGCs for novel classes of 4-alkyl-L-proline-containing metabolites and could expand the currently known three classes by over 16, as shown in the CluSeek-generated gene cluster similarity network (Fig. 3B). The network not only highlights the diversity of these newly predicted metabolite classes but also reveals their broader taxonomic distribution - most notably, identifying their presence for the first time outside of Actinomycetota, particularly in Cyanobacteriota and Pseudomonadota (e.g., class 14). Remarkably, six of the new classes (3, 6, 8-10, and 13) originate exclusively from *Streptomyces*, a genus extensively studied within Actinomycetota.

**Fig. 3.**
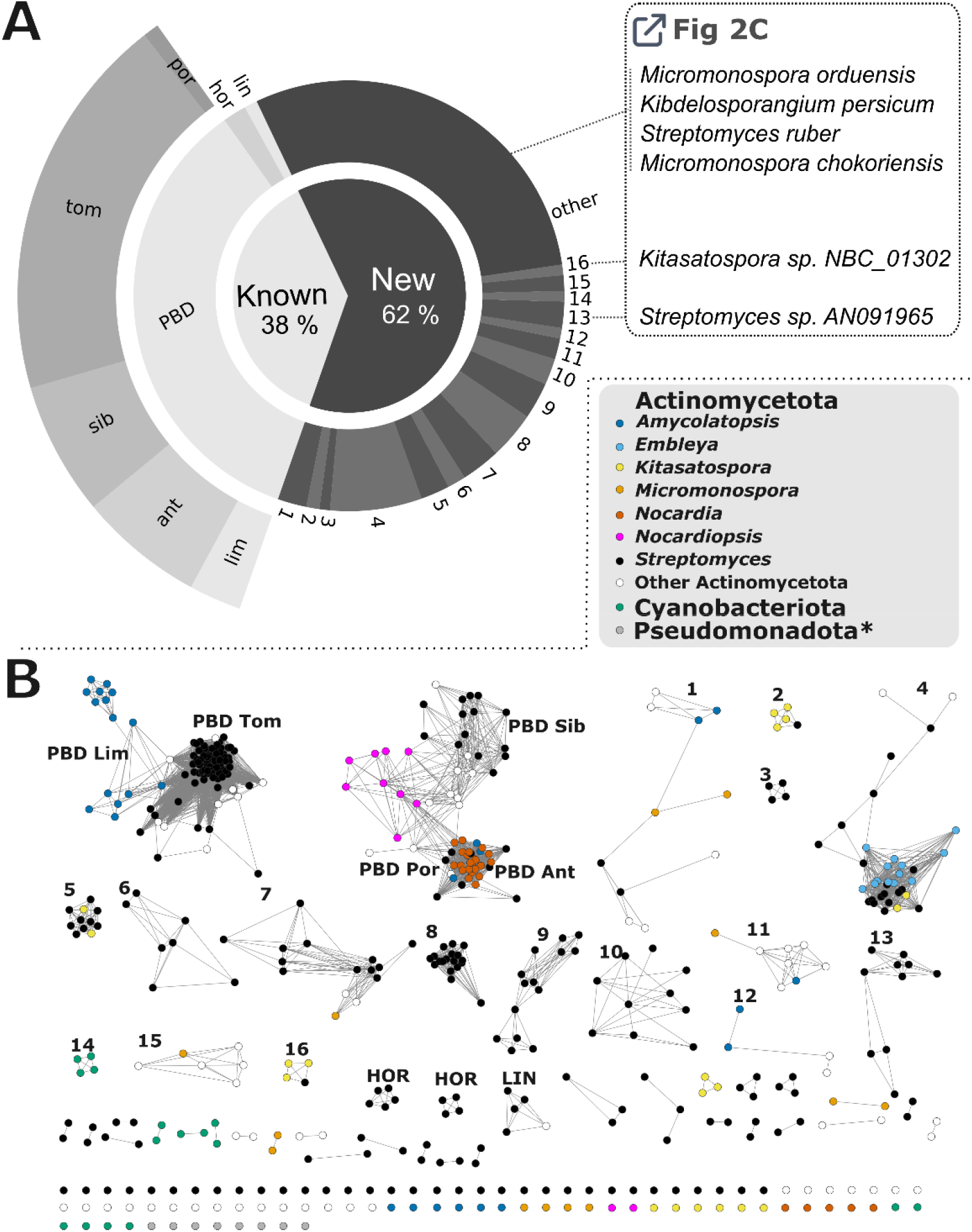
CluSeek-based genome mining of metabolites with 4-alkyl-L-proline motif. **A. Overview of gene clusters**. Known classes include lin (lincomycin), hor (hormaomycin), and PBDs (pyrrolobenzodiazepines), the latter further divided into biosynthetic and structural subclasses, lim (limazepine), tom (tomaymycin), sib (sibiromycin), por (porothramycin), ant (anthramycin). New classes are designated by numbers 1-16, each representing gene cluster type with at least 4 members; cluster types with 1-3 members are grouped as “other”. Gene clusters corresponding to six examples of new metabolites are shown in Fig. 2C. **B. Gene cluster network**. Each node represents a gene cluster; edges connect those with similar gene composition, suggesting production of structurally related metabolites. Node colors indicate the taxonomic origin. *includes also two nodes corresponding to Bacteriodota and Candidatus Dormiibacterota.

Interestingly, novel biosynthetic strategies may be discovered, as some of the 317 newly identified gene clusters do not encode nonribosomal peptide synthetase (NRPS) or any other known assembly machinery enzymes, while others contain polyketide synthase (PKS) genes. In contrast, the three previously known compound classes with the 4-alkyl-L-proline motif are assembled exclusively by NRPS (hormaomycin and pyrrolobenzodiazepines)^18,19^ or by a lincosamide-specific condensing system with NRPS-like elements (lincomycin)^32,33^, and none of them incorporate PKS-derived polyketide structures. Furthermore, a number of the gene clusters putatively encode a compound with a new variant of the 4-alkyl-L-proline motif, including differences in the length, saturation, and substitution of the 4-alkyl side chain (inferred from the *apd* gene subcluster composition), as well as tailoring of the motif (inferred from the presence of a putative proline hydroxylase-gene). Further tailoring that is unusual or unprecedented in the context of 4-alkyl-L-proline-containing compounds likely occurs via encoded putative halogenases, aryl sulfotransferases, or glycosyltransferases (compare panels B and C in Fig. 2).

To put CluSeek’s contribution into a broader context, it is important to recognize that most known 4-alkyl-L-proline-containing metabolites, belonging to the three previously described structural and biosynthetic classes, were discovered during the 1960s–1980s, the so-called Golden Era of antibiotics. This period saw the identification of many clinically used antibiotics and drug leads, largely derived from Actinomycetota. In contrast, recent decades have been marked by high rediscovery rates, leading to the widespread assumption that this phylum’s biosynthetic potential is exhausted, particularly in well-studied genera such as *Streptomyces*. However, our GenBank-scale analysis using CluSeek challenges this view. It reveals that 4-alkyl-L-proline-containing metabolites are not only more taxonomically widespread, but also significantly more structurally and biosynthetically diverse than previously recognized. Notably, several newly identified biosynthetic classes were found exclusively in *Streptomyces*, a genus often deprioritized in current discovery strategies in favor of rare Actinomycetota taxa or isolates from extreme environments. These findings challenge the prevailing approach and demonstrate that even well-characterized strains can harbor not just residual, but substantial and previously overlooked metabolic diversity. While the Golden Era yielded the first three classes via classical bioassay-guided discovery, our study suggests that the next wave, comprising 16 or more new classes, awaits discovery in the post-2025 era via targeted genome mining approach with CluSeek playing its role.

### Case study 2 - Bacterial secretion systems

Bacteria utilize multiprotein assemblies known as secretion systems to transport molecules across the bacterial cell envelope, enabling them to colonize and survive in diverse environments. A remarkable example of these systems is the type III secretion system (T3SS), which allows for the injection of type III effector proteins directly from the bacterial cytosol into the host cell cytoplasm, where they modulate host cell processes. The T3SS is widely distributed in Gram-negative bacteria and central to the pathogenesis of species, such as *Salmonella, Shigella, Yersinia*, enteropathogenic *Escherichia coli* (EPEC), and *Bordetella*^34^. The more than 20 components of the T3SS apparatus, referred to as the injectisome, are usually encoded in a gene cluster located on a pathogenicity island in the bacterial chromosome or on a plasmid, and have been found to be horizontally transferred between bacterial species^2,35^. The evolutionary distinct bacterial species may have phylogenetically related injectisomes and vice versa^36,37^. In this case study, we used CluSeek to investigate the *Bordetella* secretion complex (Bsc), which is an orphan T3SS that does not clearly belong to any of the major T3SS families^38^. As markers for this CluSeek analysis of GenBank data, we used the proteins of the inner and outer membrane rings, BscJ and BscC, respectively, the gatekeeper protein BopN, the major translocator protein BopB, and the tip filament protein Bsp22.

CluSeek identified and visualized 2228 gene clusters containing these markers (Supplementary Fig. S2). Consistent with previous reports^39,40^, cluster was present in the genomes of classical *Bordetella* species, *B. pertussis, B. bronchiseptica*, and *B. parapertussis*, which infect the respiratory tract of mammals, as well as in non-classical *Bordetella* species, *B. ansorpii* (Fig. 4A). However, our analysis also revealed that the taxonomic distribution of the Bsc-related clusters in the order Burkholderiales, and specifically in the families Alcaligenaceae and Comamonadaceae, is substantially broader than previously recognized. Within Alcaligenaceae, the Bsc cluster is common in the genus *Achromobacter*, including *A. xylosoxidans, A. insuavis, A. ruhlandii, A. dolens, A. denitrificans* and *A. deleyi*, and has also been found in the genus *Castellaniella*, including *C. ginsengisoli* (Supplementary Fig. S2 with representative examples in Fig. 4A). While *Achromobacter* species are environmental bacteria whose clinical relevance is increasingly recognized, especially in immunocompromised individuals and patients with cystic fibrosis^41,42^, *C. ginsengisoli* has only recently been associated with disease in poultry^43^. In contrast, Bcs clusters are less common within Comamonadaceae, though they were still found in *Acidovorax, Comamonas*, and *Variovorax*. Outside these families, Bcs-related clusters were only rarely detected.

**Fig. 4.**
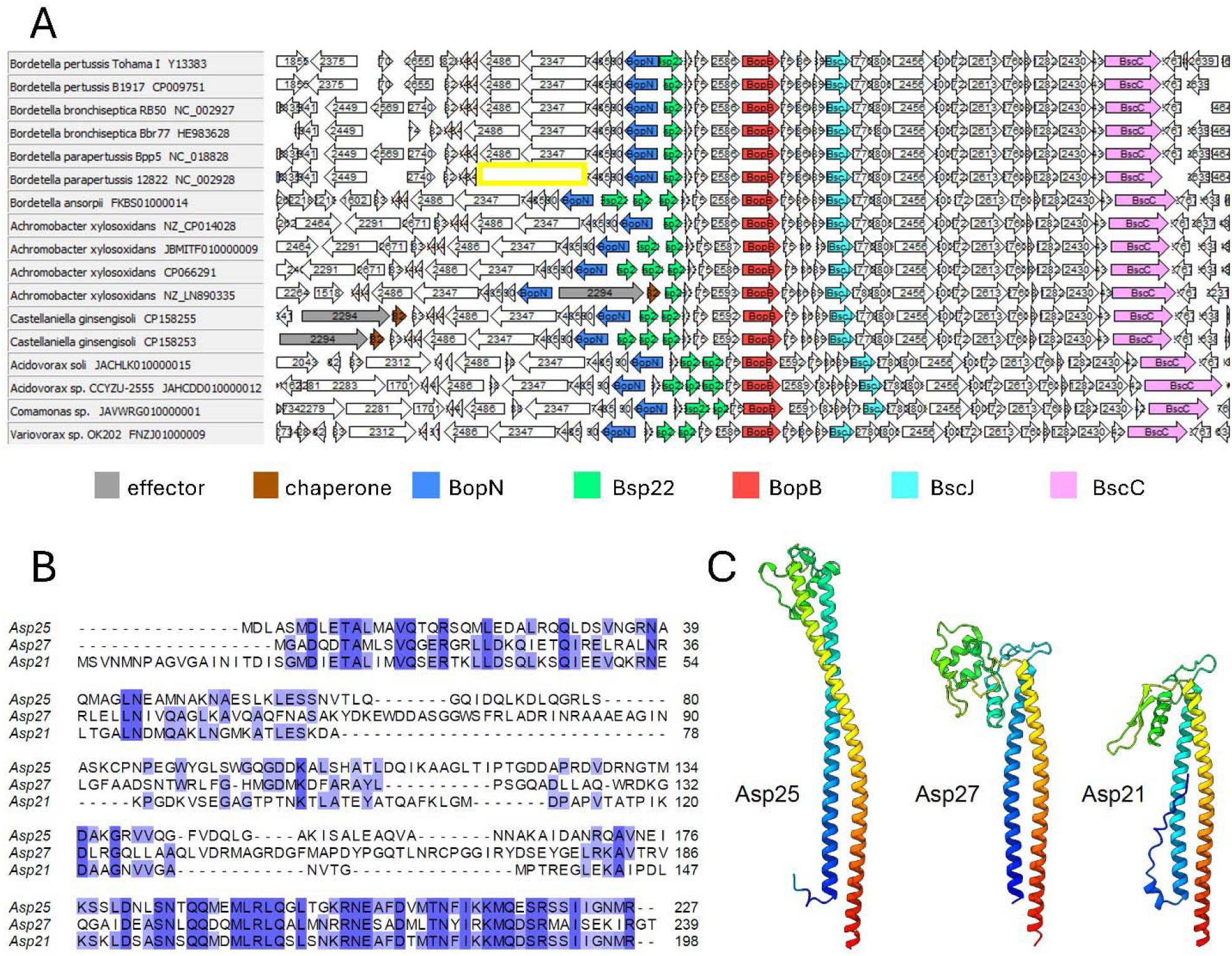
CluSeek-based analysis of the Bsc cluster encoding type III secretion system. **A. Examples of identified gene clusters in various species.** Pseudogenization of the T3SS gene cluster in *B. parapertussis* 12822 - loss of two genes is in yellow frame; note that the presented gene clusters differ in the number of *bsp22* and in position of *bopN* and *bscC*; the complete CluSeek output is depicted in Supplementary Fig. S2; gene cluster borders are not indicated. **B. Sequence alignment of Bsp22-like proteins of *A. xylosoxidans* FDAARGOS_984**. Amino acid sequences were aligned using the Clustal Omega online alignment tool available at the HHpred website (https://toolkit.tuebingen.mpg.de/tools/hhpred) and visualized with the percentage identity scheme in Jalview. Asp21 contains an apparent N-terminal extension of 20 amino acids based on the NCBI sequence, although it is plausible that the true start codon is the internal methionine (M21), which would eliminate this extension. **C. Ribbon representation of the AlphaFold-predicted monomeric structure of Bsp22-like proteins of *A. xylosoxidans* FDAARGOS_984**. This representation was generated by UCSF ChimeraX.

Regarding organization and gene content of Bsc cluster, the identified clusters are generally more uniform than the biosynthetic gene clusters analyzed in Case study 1. Nevertheless, CluSeek detected several subtle yet potentially important differences that might be otherwise overlooked. For example, while pseudogenization of the T3SS cluster in the human-adapted *B. parapertussis* 12822 (Fig. 4A) has been previously reported^44^, we identified distinct differences in Bsc-related clusters of *Achromobacter* species – an observation that was not readily expected. Specifically, a subset of *Achromobacter* strains, including *A. xylosoxidans* and *A. marplatensis*, harbored two additional genes located between those encoding BopN and Bsp22 homologs. These genes were also present upstream of the Bsc-related cluster in *Castellaniella*.

A BLASTp analysis revealed homology to an T3SS effector, the inositol phosphate phosphatase IpgD of *Shigella flexneri*, and to a T3SS chaperone, respectively (Fig. 4A). To the best of our knowledge, this putative effector–chaperone pair has not been previously linked to Bsc-related clusters or to members of these genera.

Another observation provided by CluSeek was the presence of two or even three copies of the *bsp22* gene in some strains, including *B. ansorpii* and *Achromobacter* (examples highlighted in green in Fig. 4A). Bsp22 is a structural component of the tip of the T3SS injectisome. Previous studies showed that Bsp22 of *B. bronchiseptica* RB50 is highly immunogenic, and that anti-Bsp22 antibodies can protect mice against *Bordetella* challenge^45^. This raises the hypothesis that *bsp22* gene multiplication may represent a bacterial strategy to evade host immune responses. To investigate this, we analyzed three Bsp22 homologs of *A. xylosoxidans* FDAARGOS_984 (genome accession CP066291, Fig. 4A), a clinical isolate from the sputum of a 29-year-old woman with cystic fibrosis. We designated these variants Asp25, Asp27 and Asp21 based on their approximate molecular weights (25, 27 and 21 kDa), where “Asp” stands for *Achromobacter* secreted protein. Amino acid alignments (Fig. 4B) revealed highly conserved N- and C-terminal regions that flank the variable central portion. AlphaFold3^46^ structure predictions (Fig. 4C) indicate that the conserved N- and C-terminal regions form terminal α-helices connected by a variable mixed α/β-core. In many bacteria, tip proteins form a homopentameric cap at the distal end of the needle, as in *Salmonella* and other species^47^. In contrast, in *B. bronchiseptica* and enteropathogenic *Escherichia coli* (EPEC), the needle is extended by an additional filamentous structure, formed by polymerized tip proteins Bsp22^46,48^ and EspA^49,50^, respectively. By analogy to EspA, whose helical filament structure has been resolved by cryo-EM^49,50^, the conserved terminal regions of Asp proteins may mediate filament formation, while the variable central domain, likely exposed on the filament surface, could influence antigenic properties. However, structural data for Bsp22 and/or Asp filaments are unavailable. Importantly, large-scale analysis of Bsp22/Asp homologs by Cluseek confirms that *bsp22* gene multiplication is not an artifact. CluSeek’s workflow involves a two-step protein grouping: first, by sequence similarity into subgroups; second, these subgroups are merged into broader groups through community detection on local alignment-based networks. The broader groups are used by default for gene cluster visualization, but users can also inspect the intermediate subgroups. This capability allowed us to assess amino acid divergence among Bsp22/Asp homologs. Using CluSeek’s protein group network feature, we found that these homologs predominantly fall into three distinct subgroups (Supplementary Fig. S3). Combining this with customizable filtering, we obtained a detailed view of *bsp22/asp* copy number variation across taxa, visualized at the gene cluster level (Supplementary Fig. S4), and in gene cluster networks (Supplementary Fig. S5). The analysis reveals a non-random distribution of Bsp22/Asp homologs, with gene clusters containing multiple copies typically including members from different subgroups. This conserved pattern across taxa suggests that the sequence-variable central region of Bsp22/Asp is functionally important and that maintaining multiple variants provides a selective advantage. CluSeek thus offers a robust foundation for future studies into the biological role of this phenomenon, including whether bacteria differentially express *bsp22* or *asp* gene copies, or co-express multiple variants to form heteropolymeric tip filaments with functional or antigenic advantages.

### Novelty of CluSeek

CluSeek integrates three key components: 1) identification of colocalized genes in GenBank– deposited genomes, 2) sequence-based analytical functions, and 3) visualization of the results.

Colocalization has, at least to some extent, precedent in tools like MultiGeneBlast^51^ and CAGECAT^52^, and CluSeek matches these tools in speed and result quality (see Supplementary table S1).

The analytical functions are where CluSeek truly stands out. They enable systematic, sequence-based analysis of all encoded proteins within retrieved gene clusters - a challenging task, since homology depends not only on sequence similarity but also on coverage and contextual relationships. CluSeek tackles this with a two-step approach combining sequence-similarity grouping with network-based refinement, allowing identification of homologous proteins across diverse gene clusters, including biosynthetic and secretion system loci. This, in turn, supports flexible downstream analyses and filtering options.

The third component, visualization, complements the analytical framework. Unlike CAGECAT, which is limited to 50 gene clusters and optimized for small-scale figures, CluSeek can efficiently display over gene 1000 clusters (2228 in our case study 2), supporting large-scale, unbiased analyses (see Supplementary table S1). For publication, CluSeek remains compatible with clinker for users who prefer its output.

Neither of the two case studies - representing distinct microbial scenarios - could have been completed using existing tools. Without CluSeek, the conclusions would have required extensive manual curation, highlighting its unique ability to enable efficient discovery.

## MATERIALS AND METHODS

### Software Dependencies

CluSeek is implemented in Python (version 3.9.*). It employs the SQLite3 module from the Python standard library for database management, and it also uses the following third-party modules: Biopython v.1.79 for parsing of BLASTp output and Entrez network functions^53^, PySide2 (bindings for the Qt framework v.5.15.2; https://doc.qt.io) for GUI implementation, Openpyxl v.3.0.10 for generating .xlsx files (openpyxl.readthedocs.io), Matplotlib v.3.5.1 for the creation of graphs and data visualization^54^, and NetworkX v.3.2.1 for the visualization and analysis of protein group similarity networks^55^. All third-party packages were obtained via the Python Package Index.

In addition, CluSeek relies on non-Python dependencies, namely NCBI BLASTp for remotely searching the NCBI database^10^, and the DIAMOND software suite for clustering and local alignment of protein sequences^12,56^. The software is dependent on the online database resources provided by NCBI (e.g. nuccore) ^11^ and programmatic access to them via the Entrez interface^57^.

### Project Files

CluSeek stores data in project files implemented as SQLite3 databases with a custom extension. During runtime, the database is kept in memory and is copied to the hard drive when the user saves the project. Each project file retains all information downloaded from NCBI including BLASTp search results, downstream analyses such as protein grouping and filtering, and any user modifications.

### BLASTp Search to Identify Sequence Homologs of Markers

After the sequences of markers are entered (either as FASTA or by proxy as NCBI accession codes), CluSeek accesses the online BLASTp API via Biopython’s NCBIWWW module and runs a search against the nr database. Once the NCBI servers complete the requested BLASTp search, the results are downloaded as an XML file and processed. For convenience, this query is submitted via CluSeek’s GUI, although users may also import the XML-formatted output of a previously completed BLASTp search, such as one generated via NCBI’s web interface. Note that when using the NCBI web interface, the nr database must be selected, as the ClusteredNR database is not compatible with CluSeek.

### Colocalization of Marker Homologs

After the accession codes of all relevant marker homologs are extracted from the BLASTp output, CluSeek queries the NCBI IPG resource (via Biopython/Entrez) to retrieve (i) all proteins identical to the identified homologs (including those not returned by BLASTp when searching the nr database) and (ii) the genomic position of each protein’s coding sequence. Using this positional data, CluSeek scans all contigs containing at least one marker to identify genomic regions which meet the user-defined criteria for a gene cluster. These criteria are based on (i) the presence or absence of the specific markers, and (ii) the *maximum gene cluster size* defined as the distance between the outermost markers (60 kbp by default). To provide flexibility, CluSeek applies a scoring system in which each marker is assigned a value that is awarded when its homolog is detected within the region. The cumulative score must exceed a threshold for the region to qualify as a gene cluster (by default, this threshold equals the number of markers selected for colocalization). Overlapping gene clusters are always merged, even if this exceeds *maximum gene cluster size* setting. It is possible for the same protein coding sequence to be homologous to multiple markers and be awarded score for both.

Because unfiltered results often contain duplicates from multiple whole-genome sequencing runs, CluSeek can dereplicate output to display only one representative per strain, species, or genus, depending on user preference. If multiple gene clusters per genome are present, CluSeek by default identifies contigs originating from the same sequencing run (based on accession codes) and includes all gene clusters from that run rather than only the first. This option can be disabled in the *Markers and Colocalization* tab. For sequences with unknown or unparseable taxonomy, CluSeek treats them as belonging to separate taxonomic groups to avoid discarding potentially relevant results. Alternatively, users may enforce stricter dereplication by excluding all sequences with ambiguous taxonomy, ensuring at most one representative per group.

### Gene Cluster Visualization and Protein Grouping

To display gene clusters that meet user-defined criteria, CluSeek queries the NCBI database via Biopython/Entrez and retrieves the corresponding GenBank records. Each visualized cluster includes the outermost markers, the intervening sequence, and flanking regions of user-defined size (75 kbp upstream and downstream of the outermost markers by default, or up to the end of the contig). For downstream analysis, proteins encoded within the visualized gene clusters are grouped by sequence similarity using DIAMOND and NetworkX. First, DIAMOND’s centroid-based *cluster* algorithm groups proteins into relatively homogeneous subgroups. Then, a local alignment of all subgroup centroids against all other subgroup centroids is performed using DIAMOND’s *blastp* algorithm, generating an all-vs-all similarity matrix. This matrix is then converted into a network graph in NetworkX, and protein subgroups are further assigned to higher-order groups using NetworkX’s implementation of the Clauset–Newman–Moore hierarchical agglomeration algorithm^58^ (Supplementary Fig. S6). This algorithm maximizes modularity, ensuring that subgroups within a group are more closely related to each other than to subgroups outside the group.

### Ordering Gene Clusters

The visualized gene clusters are automatically ordered according to the Jaccard similarity of their protein group composition:

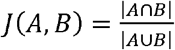

Where A and B are sets of protein groups represented in each gene cluster. Thus, gene clusters are regarded as more similar if there are more protein groups which are represented in both, disregarding the number or position of the protein group members.

### Case Study Data Analysis

The markers used in this study and their accession numbers were: Apd1 - ABX00599.1, Apd2 - ABX00598.1, BopN - WP_010930827.1, Bsp22 - WP_003820049.1, BopB - WP_010930824.1, BscJ - WP_010930821.1, BscC - ETA64901.1. BLASTp searches were performed on the NCBI website (April 10, 2025, for case study 1; April 19, 2025, for case study 2) using default parameters, except that *max target sequences* return limit was set to 5000 for both case studies, and dereplication to strain level was applied in case study 2. The resulting XML files were imported into CluSeek. For downstream analyses, we used CluSeek 2.0.1 with default settings except for the *maximum gene cluster size* parameter, which was reduced from the default 60 kbp to 35 kbp in case study 2.

For gene cluster network analysis in case study 1, candidate biosynthetic gene cluster borders were estimated manually based on the presence of typical primary metabolic genes. The central regions of the clusters were then aligned and a 30 kbp central region was exported as a .gml file using a cutoff threshold of 0.55.

For case study 2, variants of the *bsp22* gene were detected and visualized using CluSeek’s protein group network analysis (accessible via *Info* sub-tab). For the gene cluster network analysis, the gene clusters were exported as a .gml file with a cutoff threshold of 0.95. The borders of the putative T3SS gene clusters were defined by the *cesT* family type III secretion chaperone gene at one end and the *bscC* gene at the other. For the gene cluster network analysis, the gene clusters were exported as a .gml file with the cutoff threshold 0.95.

The gene cluster and protein group networks were visualized in Cytoscape.

### CluSeek Benchmarking Against CAGECAT

The benchmarking focused on identifying gene clusters containing colocalized Apd1 and Apd2 markers from case study 1. Search criteria were matched between the tools: BLASTp searches were run with an e-value threshold of 10□^2^ and no identity or coverage thresholds. Loci where both Apd1 and Apd2 were identified within a maximum intergenic distance of 60 000 bp were considered hits. Time measurements were performed in triplicate, with each replicate initiated simultaneously to minimize bias from variable NCBI server load.

The first replicate was used to assess qualitative differences in the results. Accession codes of all hits were extracted to compare sequences identified by both tools versus those found uniquely by one. Because sequences may be mirrored between the RefSeq and nuccore databases^59^ (distinguished by accession code prefixes), CAGECAT^52^ appears to apply dereplication by preferentially selecting RefSeq entries when available. To achieve parity, a manual dereplication step was applied to CluSeek results (targeting sequences with an underscore as the third character in the accession code), as CluSeek was run with the *Do Not Dereplicate* setting.

Gene cluster visualization was compared purely in terms of speed due to the large qualitative differences in the outputs. For this assessment, gene clusters containing Apd1 and Apd2 obtained from the previous search were used. As the version of clinker available via CAGECAT cannot display more than 50 gene clusters at the same time, a randomized set of 10 and 50 gene clusters was used for this purpose. The clusters were first selected and visualized in CluSeek, then exported into CAGECAT. The 10-gene cluster test was run in triplicate, while the 50-gene cluster test was performed only once due to its long run time and apparent strain on host server.

The reference values for the maximum number of gene clusters displayed for CAGECAT and CluSeek were based on the CAGECAT software limit, and user-reported observations during testing, respectively.

## CONCLUSION

CluSeek offers a fundamentally different approach to genome mining compared to widely used tools such as antiSMASH. Instead of relying on predefined rules or curated libraries of known gene clusters, CluSeek uses protein sequences to scan the entire GenBank database and identify gene clusters encoding two or more user-selected marker proteins. Its intuitive graphical user interface and interactive outputs make CluSeek a uniquely accessible and customizable tool for genome mining. It follows a simple download–open–use principle and requires no complex installation. CluSeek is freely available at https://cluseek.com.

Our case studies demonstrate the broad applicability of CluSeek across different biological contexts. In the first case study, we used CluSeek to identify novel biosynthetic gene clusters encoding 4-alkyl-L-proline-containing metabolites in Gram-positive bacteria, potentially expanding the number of known metabolite families from three to more than sixteen based on GenBank data. In the second case study, we applied CluSeek to explore type III secretion systems in Gram-negative bacteria, where it uncovered pseudogenization events, previously unreported T3SS effector-chaperone pair, and notable gene multiplications. These findings allowed us to formulate a hypothesis on their biological significance, supported by CluSeek’s ability to perform GenBank-scale analyses.

CluSeek enables the discovery of biosynthetic gene clusters encoding metabolites with shared structural features, even when they do not belong to well-characterized NRPS, PKS, or RiPP families, something conventional genome mining tools are not designed for. More broadly, it can be applied to explore any gene cluster or any phenomenon involving at least two colocalized genes, from global-scale analyses of gene cluster composition to detecting subtle variations and examining the taxonomic distribution of the phenomenon.

## DATA AVAILABILITY

Supplementary tables and figures are included in the Supplementary Information. Supplementary Figures S1, S2, and S4, along with the .clp files of CluSeek projects from the two case studies, and .cys files with gene cluster and protein group networks are available on Zenodo https://doi.org/10.5281/zenodo.17062728. Tutorial videos are available on YouTube at https://www.youtube.com/@CluSeek-p6u.

## CODE AVAILABILITY

CluSeek is available for Windows and MacOS as a standalone executable, or as a Python package named `cluseek` available from the Python Package Index for Windows, MacOS and Linux. All versions are packaged with a Diamond sequence alignment software executable which for Windows users necessitates the installation of Visual C++ x86 redistributables. Visit https://cluseek.com to download the latest version of CluSeek, complete with a quick installation guide. This publication describes CluSeek 2.0.1, the source code of which has been made available on GitHub (https://github.com/HrebicekO/CluSeek) and Zenodo (https://doi.org/10.5281/zenodo.17062728).

## ACKNOWLEDGEMENTS

This work was supported by the projects National Institute of virology and bacteriology (Programme EXCELES, ID Project No. LX22NPO5103) - Funded by the European Union - Next Generation EU, Talking microbes - understanding microbial interactions within One Health framework, supported by the Ministry of Education, Youth and Sports of the Czech Republic grant No. CZ.02.01.01/00/22_008/0004597 and grants 25-16251S to ZK and 24-11053S to JK of the Czech Science Foundation.

