## Supplementary Information for "CluSeek: Bioinformatics Tool to Identify and Analyze Gene Clusters"

**SUPPLEMENTARY TABLES**

**Supplementary Table S1** Benchmarking of CluSeek against CAGECAT.

|  |  | CluSeek | CAGECAT |
| --- | --- | --- | --- |
| Colocalization (Apd1, Apd2) | Average time (n=3, s) | 434 | 444 |
|  | Unique accession codes detected | 788 | 775 |
|  | Accession codes only found by one and not the other | 14 | 1 |
| Visualization (Apd1, Apd2 gene clusters) | Gene cluster display limit | > 1 000 | 50 |
|  | Average time (n=3, s) to display 10 gene clusters | 18 | 275 |
|  | Time (n=1, s) to display 50 gene clusters | 78 | 13 587 |
|  | *Time to display all results not available as it exceeds CAGECAT's display limit.* | | |

In numerical terms, CluSeek and CAGECAT perform similarly in colocalization, both in terms of speed and the number of results yielded. The key difference was observed in the visualization step. Qualitative differences aside, CluSeek can display significantly more gene clusters and and does so in markedly less time. Additional features of CluSeek, such as protein grouping, gene cluster and protein group networks, and advanced filtering, cannot be benchmarked against CAGECAT, as they are not available in that tool.

**SUPPLEMENTARY FIGURES**

Supplementary Figures S1, S2 and S4 are available as separate files in high resolution and are also accessible at Zenodo (<https://doi.org/10.5281/zenodo.17062728>). Outputs corresponding Supplemetary figures S1 and S2 are available as .clp also on ZENODO and these can be open and inspected in CluSeek.

**Supplementary Figure S1** Gene clusters identified within GenBank using CluSeek and Apd1 and Apd2 markers; *apd1-apd6* of the *apd* subcluster encoding the biosynthesis of the 4-alkyl-L-proline motif are in red; biosynthetic gene cluster borders are not indicated. The figure is available on Zenodo (<https://doi.org/10.5281/zenodo.17062728>).

**Supplementary Figure S2** Gene clusters identified within GenBank using CluSeek and markers of the type III secretion system; Homologs of BopN - blue, Bsp22 - green, BopB - light red, BscJ - light blue, BscC - pink; secretion system gene cluster borders are not indicated. The figure is availbale on Zenodo (<https://doi.org/10.5281/zenodo.17062728>).

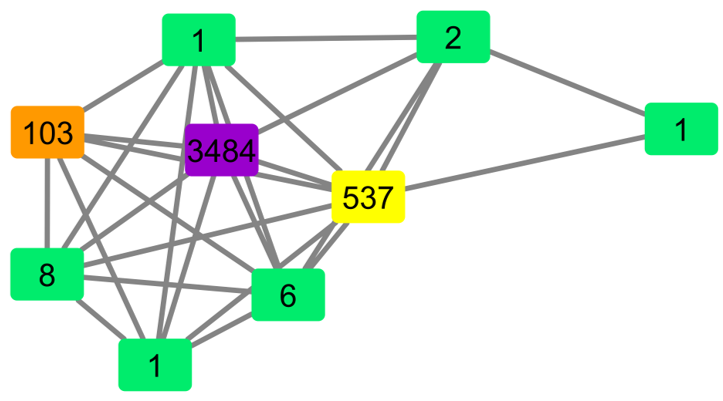

**Supplementary Figure S3** Protein subgroups within homologs of Bsp22. Numbers indicate the number of members in each subgroup. The three main subgroups are highlighted in yellow, orange, and violet. Remaining subgroups retain the original green color used for all Bsp22 homologs in Supplementary Figure S2.

**Supplementary Figure S4.** Modified version of Supplementary Figure S2, with Bsp22 homologs colored according to their subgroup assignments. Colors correspond to those shown in Supplementary Figure S3. The figure is available on Zenodo (<https://doi.org/10.5281/zenodo.17062728>).

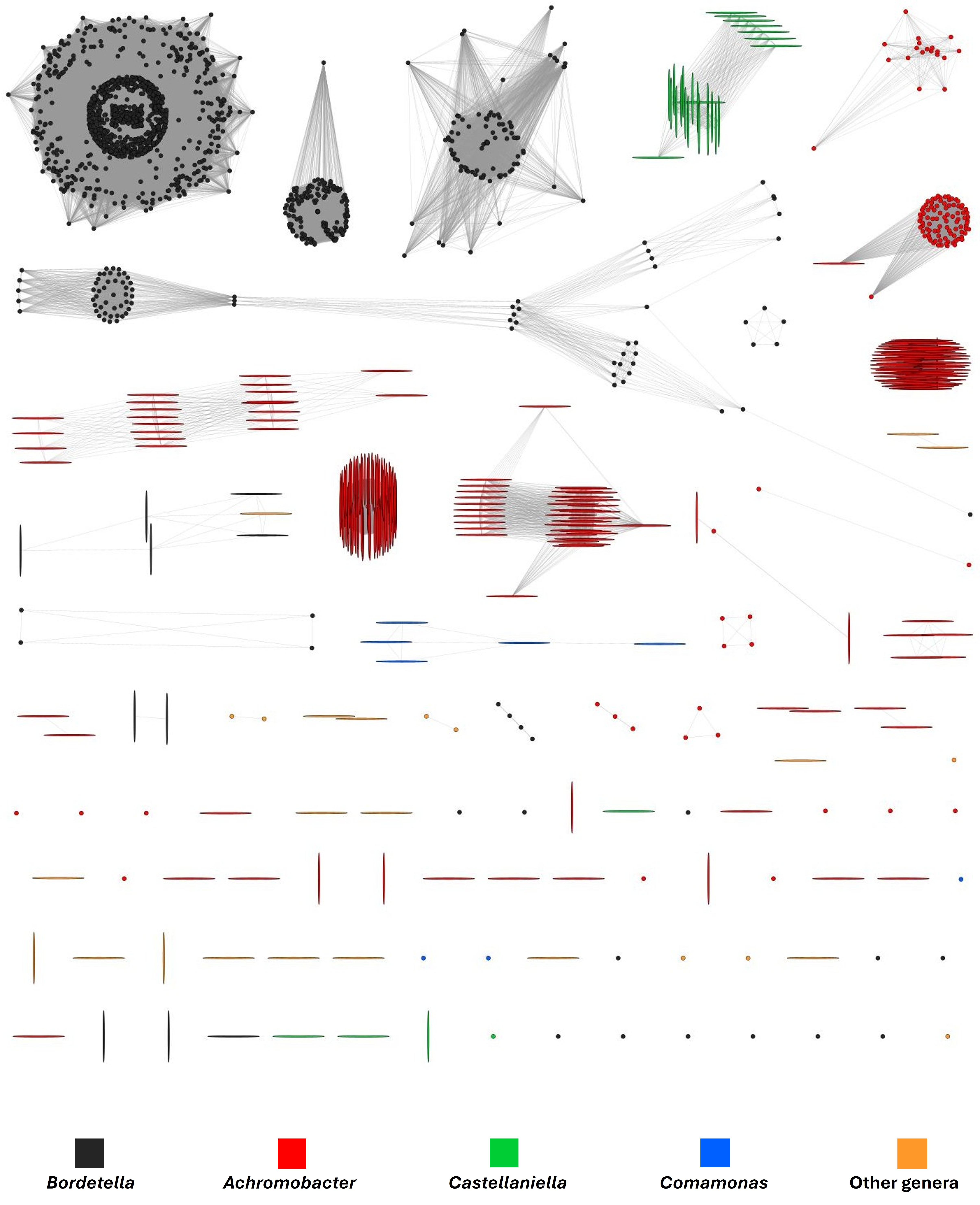

**Supplementary Figure S5** Gene cluster similarity network of T3SS gene clusters from case study 2. Number of homologs of *bsp22* in a gene cluster: dot – 1 homolog, horizontal bar – 2 homologs, vertical bar – 3 homologs.

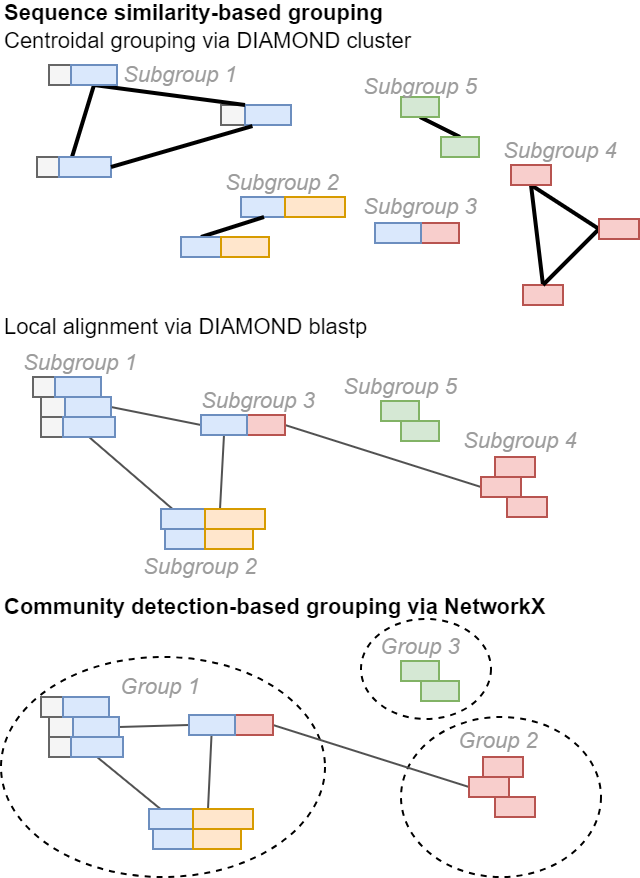
**Supplementary figure S6:** Protein grouping algorithm. Protein similarity is precalculated at two levels. Higher-homogeneity subgroups are created using DIAMOND’s sequence-similarity-based centroid clustering. Pairwise similarities are further refined by local alignment (DIAMOND blastp), and final protein groups that are applied in CluSeek´s output are assembled through community detection–based clustering in networkX.
