## Supplementary figures and images for "CluSeek: Bioinformatics Tool to Identify and Analyze Gene Clusters"

### Supplementary Figure S1

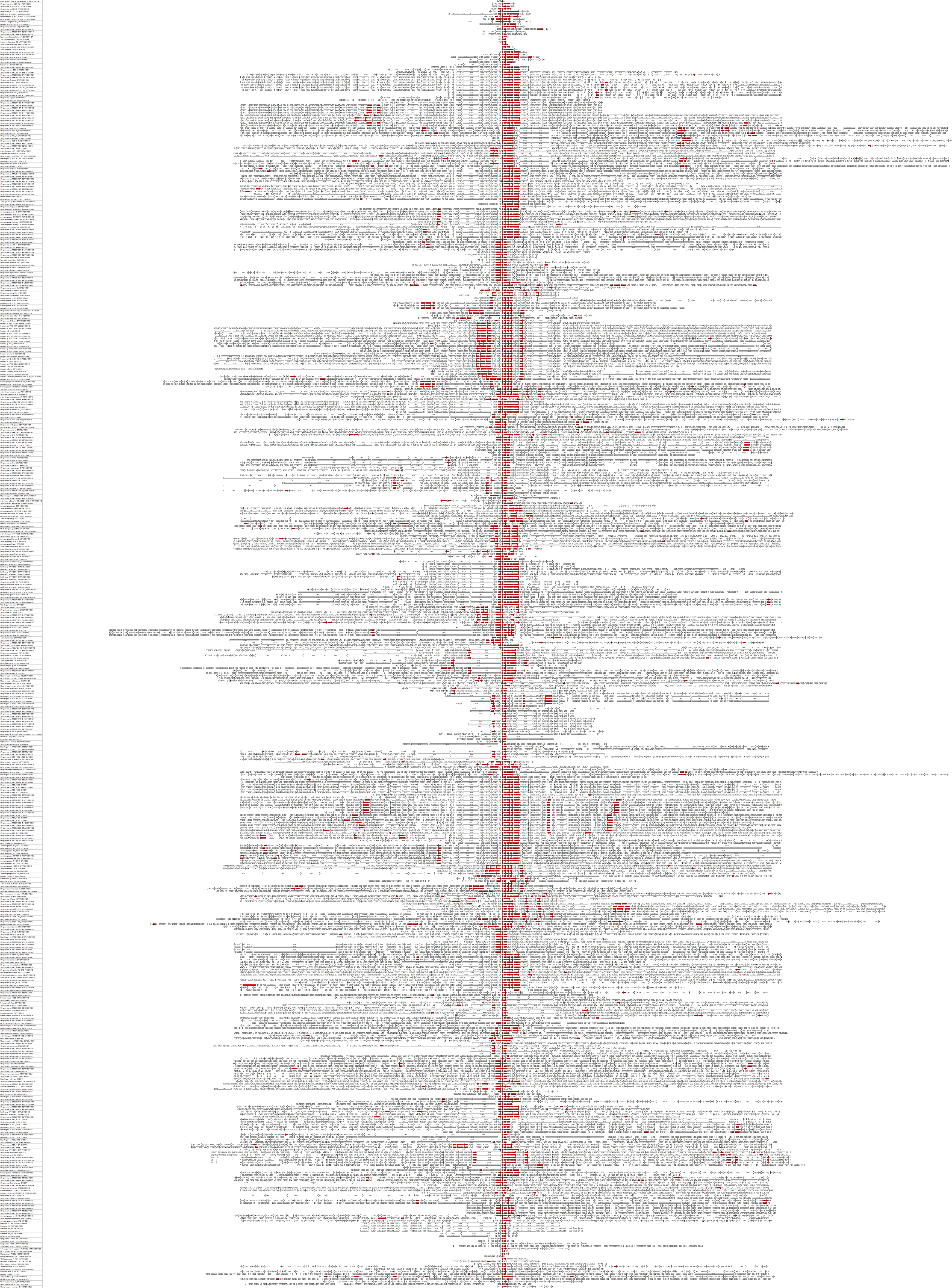
